# Geometrically balanced model of cell growth

**DOI:** 10.1101/2024.11.04.621929

**Authors:** Alexei Vazquez, Tomáš Gedeon

**Affiliations:** Nodes & Links Ltd, Salisbury House, Station Road, Cambridge CB1 2LA, UK; Department of Mathematical Sciences, Montana State University, Bozeman, MT USA

## Abstract

The proteome balance constraint in metabolic flux balance analysis asserts that the proteome is constructed by ribosomes, which themselves contain many proteins. This leads to a fundamental question of optimal allocation of limited proteome among different pools of enzymes, which include ribosomes themselves. However, recent work points to additional constraints imposed by the cell geometry. In this paper we deduce the *proteogeometric constraint* 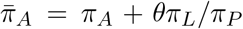, where *π*_*A*_, *π*_*P*_ and *π*_*L*_ are the proteomic fractions allocated to the cell surface area, protein synthesis and cell membrane phospholipids synthesis and 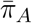 and *θ* are constants imposed by geometry of the cell. We illustrate the relevance of this constraint using a reduced model of cell metabolism, illuminating the interplay between cell metabolism and cell geometry.

## 1 Introduction

In the last 20 years we have advanced our understanding of physical and chemical constraints shaping the metabolic capabilities of cells. It all started with mass balance, underpinning the creation of flux balance models [1, 2, 3, 4, 5, 6, 7, 8]. It was followed by molecular crowding of the cell volume [9, 10, 11], the proteome balance constraint [12] and molecular crowding of the cell membrane [13]. Today the proteomic balance constraint is well stablished and used in most studies [14, 15, 16, 17, 18, 19, 20, 21]. However, recent publications have highlighted the need for a deeper understanding of the mode of operation of the crowding constraints of the cell volume and the cell membrane area [22, 23, 24].

At the intuitive level, making more cell membrane implies a metabolic cost and a natural selection for increased molecular crowding [23]. At the same time, nutrient transporters and other metabolic enzymes are located in the cell membrane, which would favor larger cellular membrane, or at least imply a requirement of a minimal area of membrane that can support required growth rate [13, 25, 24]. The surface area that holds the cell membrane components also encloses the volume which contains all cell components. Finally, these allocation processes need to happen under the self-replicative nature of protein synthesis: ribosomes make new ribosomes and synthesis of any other protein takes away resources from ribosome synthesis, which is the basics of the proteomic constraint [15].

Here we derive the basic equation for the volume to area balance that is consistent with the metabolic requirements of a cell. We also analyze a reduced model focusing on interpretation. In spite of its simplicity, this reduced model predicts proteomic fractions that are close to those observed for the bacterium *E. coli* and brings new insights into the interplay between molecular crowding and the cell membrane area to volume ratio under different conditions. Our work demonstrates that the cell volume-area constraint is independent of the proteome allocation constraint and should be included in flux balance models of cell metabolism.

## 2 Volume to area balance

To derive a volume to area balance constraint we focus on major macromolecular synthesis processes. That includes protein synthesis driven by ribosomes, RNA nucleotide synthesis catalyzed by RNA polymerase and membrane synthesis by some membrane synthesis pathway. We denote by 𝒫, 𝒩 and ℒ the set of all metabolic enzymes required for the biosynthesis of proteins, RNA nucleotides and cell surface area lipids. We further denote by ℳ = {𝒫, 𝒩, ℒ, …} the full set of enzymes, including any additional enzymes that will be used in our cell metabolism model.

Our core flux balance equations reflect protein, RNA and membrane lipid balance

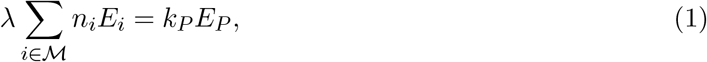

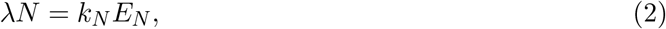

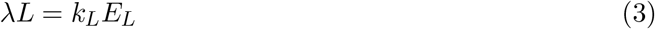

where λ is the growth rate, *E*_*i*_ is the moles of *i*-pathway enzymes per cell, *n*_*i*_ is the moles of amino acids per mole of enzymes of pathway *i, N* the number of RNA nucleotides per cell, *L* are the moles of lipid bilayer units per cell, and *k*_*i*_ is the *i*-th pathway turnover number.

These flux balance equations are complemented by the geometric balance equations for cell volume *V* and cellular surface area *A*

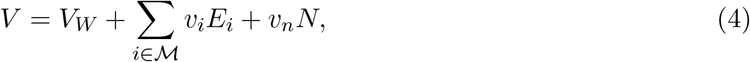

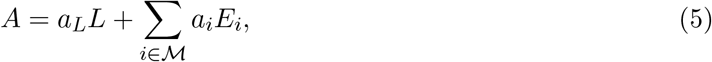

where *V*_*W*_ is the volume of water, *υ*_*i*_ the molar volume of enzyme *i, υ*_*n*_ the RNA volume per RNA nucleotide and *a*_*i*_ the molar cell membrane area occupied by enzyme *i*, in units of area per mole of enzyme. Note that *a*_*i*_ = 0 for enzymes that do not localize to the cell membrane.

We now introduce important quantities that will simplify our model. We measure the total protein content in units of number of amino acids in proteins and denote its magnitude by

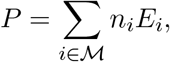

the macromolecular volume fraction by

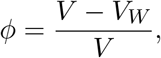

the RNA to protein ratio by

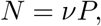

the proteomic fractions by

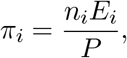

and the proteomic fraction located at the membrane by

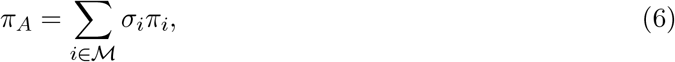

where the the subscript *A* stands for the cell area and *σ*_*i*_ is the fraction of the *i*-th protein pool that is allocated to the membrane.

We assume that the molar volumes and areas of enzymes scale with their size quantified in number of amino acids

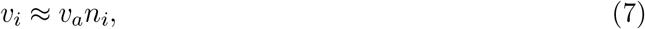

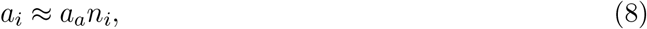

where *υ*_*a*_ and *a*_*a*_ are the typical *i*-th pool protein volume and area per amino acid. Deviations from this scaling are negligible compared to the uncertainty about the enzyme turnover numbers. The scaling approximations (7) and (8) simplify the analysis significantly. For example

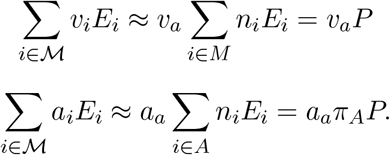

We should bear in mind these are approximations. A more precise model should take into account the variability of the specific protein volume and area across proteins. These numbers are not available in general, especially for the protein area requirements. In that case we would need to work directly with the constraints imposed by equations (4) and (5).

Finally, taking into account that

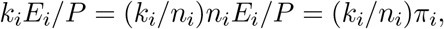

we define the specific turnover rates

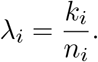

Note that we define the specific rates per unit of amino acid, because we count protein content in units of number of amino acids. That is in contrast to the more common use of specific rates per protein mass. We argue that the specific rates per unit of amino acid have better measurement units for interpretation (1/time). In fact, as illustrated below, λ_*i*_ can be interpreted as the cell growth rate if the pathway *i* would be limiting.

The equation (1) can be then rewritten as

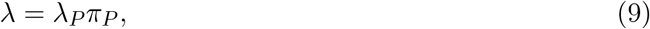

and the equation (2) as

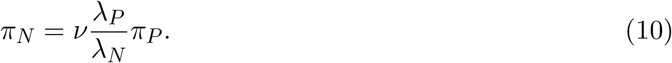

Using (9) the membrane balance equation (3) becomes

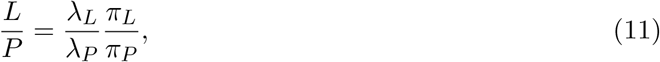

Finally, the volume balance and area balance equations (4)-(5) become

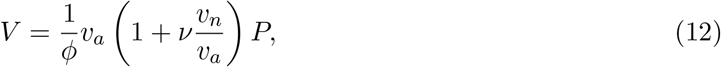

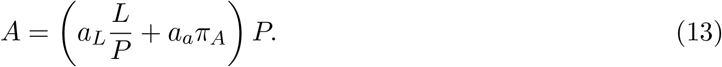

Our aim is now to obtain a membrane-volume balance equation from the equation (13). Dividing by *P* and using (11) we get

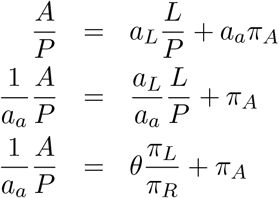

where in the last equation we introduced

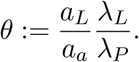

At this point we introduce a surface to volume scaling. Our cell population level model operates on time scales longer than the cell cycle. Because of this longer time scale we do not take into account geometrical changes during the cell cycle and assume that the cell’s geometry does not change. Given that the cell volume to area ratio is a key variable in bacterial morphogenesis [26], we parametrize the cell geometry by the cell volume to area ratio

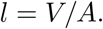

With this scaling and using (12) we obtain the final equation

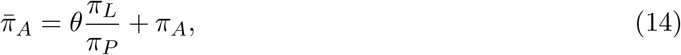

where we again introduced a new constant on the left hand side

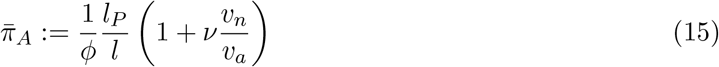

where

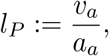

is the typical volume to area ratio of proteins.

We arrive at our final model in which the membrane-volume balance equation (14) is complemented by the growth rate equation, from (9),

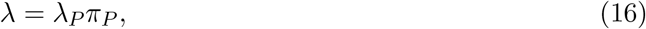

RNA polymerase proteome fraction, from equation (10),

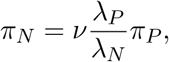

and the proteome balance

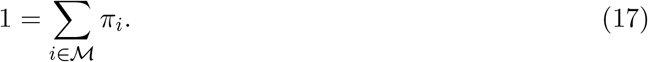

The last three equations are part of a standard proteome constraint approach. The membrane-volume balance equation (14) is the new constraint.

## 3 General observations

We call equation (14) the *proteogeometric constraint*, as it relates proteomic fractions and the cell geometry. It also serves as a self consistency check between geometry and macromolecular composition. We can reach important conclusions by inspecting this equation. First, it is obvious that the proteomic fraction at the cell membrane is bounded by the cell geometry: 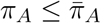. If we specify the linear dimension of proteins *l*_*P*_ and the cell *l* and the macromolecular volume fraction *ϕ*, then 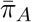 gives us the maximum proteomic fraction that can be located in the cell surface area. For example, based on that for the PTS glucose transporter of *E. coli, l*_*P*_ ∼ 0.009 *µ*m, cell volume/area measurements for *E. coli* report *l* ∼ 0.15 and the macromolecular volume fraction is about *ϕ* ∼ 0.4 (Supplementary Information), resulting in 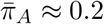. That is, only about 20% of the cell proteome of *E. coli* can be located at the cell membrane. Experimental values are scattered around 20% [27, 28], indicating that the upper bound 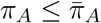 suggested by our simple model is correct.

The second observation is the role of the proteomic fraction *π*_*L*_ of the membrane lipids synthesis pathway, which enters the equation divided by the proteomic fraction of the protein synthesis pathway *π*_*P*_. The numerator *π*_*L*_ accounts for membrane lipids that are required to have a closed surface area, and the denominator *π*_*P*_ describes the bulk of proteins contained within the cell membrane area. This introduces a non-linearity, in contrast with previously used assumptions of linear constraints in the proteomic fractions. In fact, finding the optimal metabolic rates under the proteogeometric constraint becomes a quadratic optimization problem. This quadratic programming problem is convex because it contains only one non-linear term (*π*_*L*_*π*_*P*_). Therefore, its numerical solution scales linearly with the number of proteomic fractions in the model. That is relevant when applying the proteogeometric constraint to genome scale models.

Finally, the proteogeometric constraint (14), which emerged from the area constraint (5), that describes what fraction of the cell membrane is occupied by proteins or by lipids. In the right hand side, *π*_*A*_ is a scaled version of the cell membrane fraction occupied by proteins, while the remaining 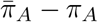 is is a scaled version of the area occupied by membrane lipids. Therefore, we can use the ratio

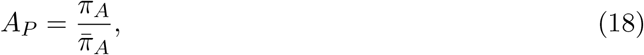

to quantify the cell membrane fraction occupied by proteins.

## 4 Simple model with energy balance

To illustrate the relevance of the volume-area balance constraint, we consider a reduced model with four enzymes/pathways: ℳ^∗^ = {ℛ, 𝒩, ℳ, ℰ}, where ℰ stands for the energy generating pathway. We further assume that amino acids and glucose are available in the extracellular media. In this way our model takes into account a trade-off between energy cost of the biosynthesis of the membrane and biosynthesis of enzymes of central metabolism and those involved in RNA synthesis. However, a further extension of our model would also acknowledge that external precursors usually enter cell through membrane based transporters and that cell needs to match the number of these transporters to the number of internal enzymes and ribosomes in a way that maximize their flux. Here we assume an existence of a pathway ℰ for energy generation with associated index *i* = *E*. First, from equation (17) we obtain the reduced protegeometric constraint

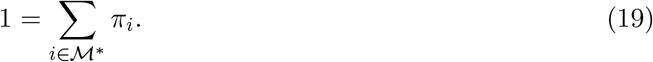

Second, from the volume-area balance equation (14) and the definition of proteomic fraction contributing to the cell membrane area (6) we obtain the reduced volume-area balance equation

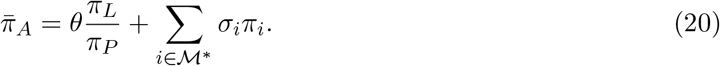

Finally, recalling that *k*_*i*_*E*_*i*_ = λ_*i*_*π*_*E*_, we add the energy balance equation

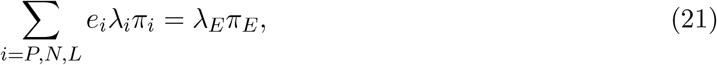

where *e*_*i*_ is the energy cost of pathway *i* in units of dephosphorylations per ligated building block.

We use equation (21) to solve for the proteomic fraction of energy generation

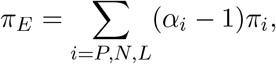

where

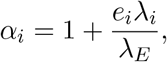

Inserting this result into (19) yields

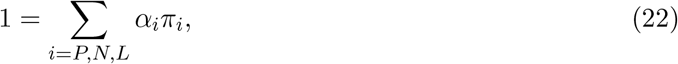

Similarly, starting with the surface balance equation (20) we get

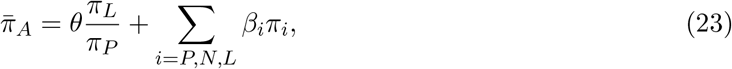

where

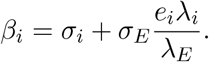

The RNA polymerase proteomic fraction is related to the the ribosome proteomic fraction via equation (10). Substituting this result into equations (22) and (23) we obtain

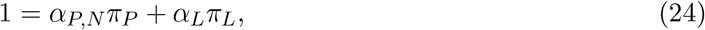

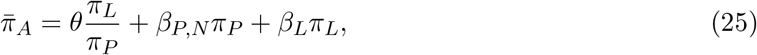

where

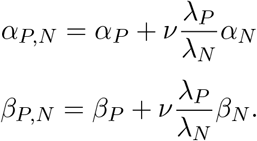

To summarize our development to this point the energy production is subject to two constraints: (a) proteome balance constraint, and (b) the proteogeometric constraint which directly led to equations (24) and (25). In our final step we solve the system of equations (24) and (25) by expressing *π*_*L*_ in terms of *π*_*P*_ from (24)

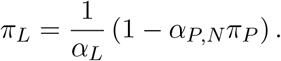

and then using (25) to arrive at our final equation

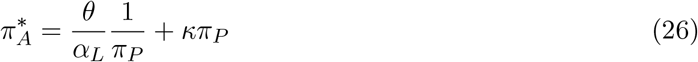

where we define new constants

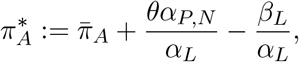

and

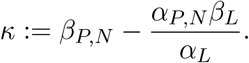

Importantly, equation (26) is quadratic equation in *π*_*P*_ and therefore has two solutions

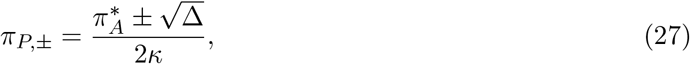

where the discriminant is defined by

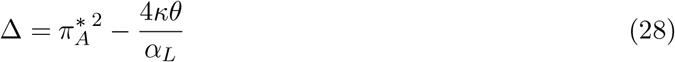

In the following we investigate whether both of these central metabolisms allocations *π*_*P*_ represent biologically meaningful solutions.

### 4.1 Parameter estimation

The key kinetic parameters λ_*i*_ quantify the enzyme (resp. pathway) rate per unit of amino acid contained in the enzyme (resp. pathway enzymes). Biologically, they quantify the cell growth rate when the corresponding enzyme or pathway is limiting. The smaller the λ_*i*_ value the higher is its importance in setting the maximum growth rate. To estimate the λ_*i*_ of a pathway we use the following reasoning.

Suppose the pathway labelled by index *i* has different enzymes *j* = 1, 2, At steady state the rates of individual enzymes *r*_*ij*_ are related to the overall pathway rate *r*_*i*_ by *r*_*ij*_ = *γ*_*ij*_*r*_*i*_, where *γ*_*ij*_ is a yield coefficient. For example, if *i* = “*glycolysis*” and we measure the glycolysis rate as the rate of pyruvate formation, then all reactions above the Glyceraldehyde 3-phosphate dehydrogenase (Gapdh) step will have *γ*_*i,j*_ = 1*/*2, while those below including Gapdh will have *γ*_*i,j*_ = 1. Then, assuming the kinetic model *r*_*ij*_ = *k*_*ij*_*E*_*ij*_ and the aforementioned relation *r*_*ij*_ = *γ*_*ij*_*r*_*i*_ we obtain

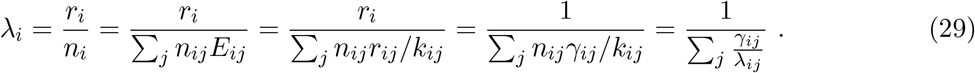

This shows that the kinetic rate λ_*i*_ of a pathway *i* is the geometric mean of the kinetic rates λ_*ij*_ for the pathway biochemical reactions. Using this approximation and known enzyme parameters (Supplementary Information) we have estimated λ_*E*_, λ_*L*_ and λ_*P*_ for *E. coli* (Table 1).

**Table 1:**
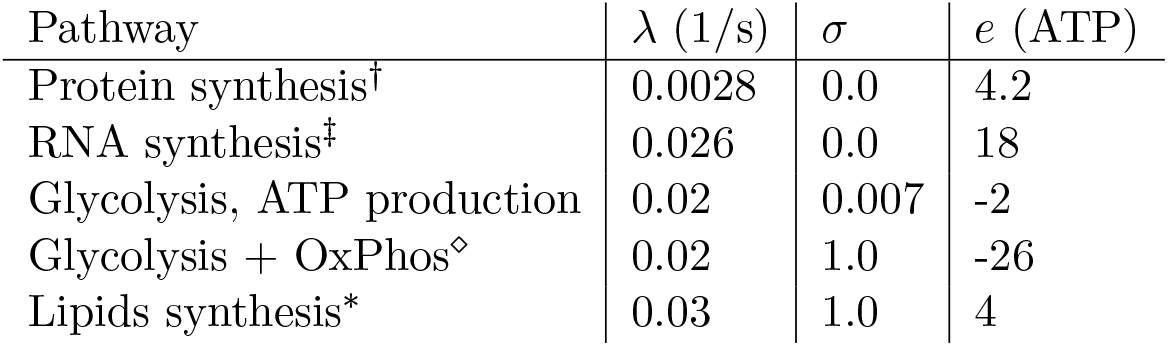
Effective parameters estimates for the reduced model pathways. ^†^Estimated from the ribosome translation rate, but neglecting the contribution of amino acids synthesis. ^‡^The λ values were estimated from the RNA polymerase transcription rate, while the *e* value takes into account the energy cost of nucleotide synthesis. ⋄ This estimate gives about the same value as glycolysis based on estimates for eukaryote cells [29]. The actual value is currently under debate [30, 31].^∗^Estimated using cardiolipin synthesis as a reference.

The specific rates λ_*i*_ are convenient to make calculations involving proteomic fractions. For example, given the pathway *i*, the proteomic sub-fraction located to the cell membrane can be calculated as

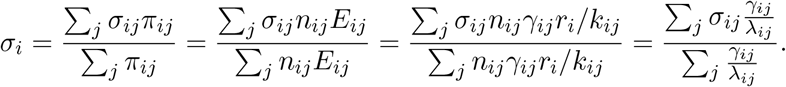

The aggregation of enzyme quantities is weighted by their inverse λ. Using this equation and and known enzyme parameters (Supplementary Information) we have estimated *σ*_*i*_ for the reduced model pathways (Table 1).

Finally, we emphasize that the kinetic parameter estimates are inaccurate and different experimental reports often yield very different values. In addition, there likely does not exist a “true value of a kinetic parameter” for a given reaction. There are isoenzymes, changes in enzyme activity due to molecular crowding or other modifying conditions, as well the dependency of the reaction rate on substrate availability. We will account for all that by utilizing random variations of the λ_*i*_ reported in Table 1. To this end we generate random numbers extracted from a triangular distribution with min, mode and max at λ_*i*_*/*2, λ_*i*_, 2λ_*i*_. Then we calculate the median and the confidence interval between quantiles 0.2 and 0.8. The goal of this simulation is to illustrate the effect of parameter uncertainty, rather than to make quantitative statements. Therefore there is no specific biological reason for the choice of distribution nor the magnitude of the variation.

### 4.2 Results

Based on our parameter estimates *κ* < 0 and the only valid solution in equation (27) is *π*_*P*,−_. Figure 1 shows the distribution of *π*_*P*,−_ solutions of equation (26), assuming that energy is generated by glucose fermentation and introducing variability in the λ_*i*_ values reported in Table 1. The mode is around 0.5, slightly higher than the experimentally observed maximum values of the ribosome proteome fraction (∼ 0.4) [32]). Yet, given the simplicity of the reduced model the close fit to the experimental data is a remarkable achievement.

**Figure 1:**
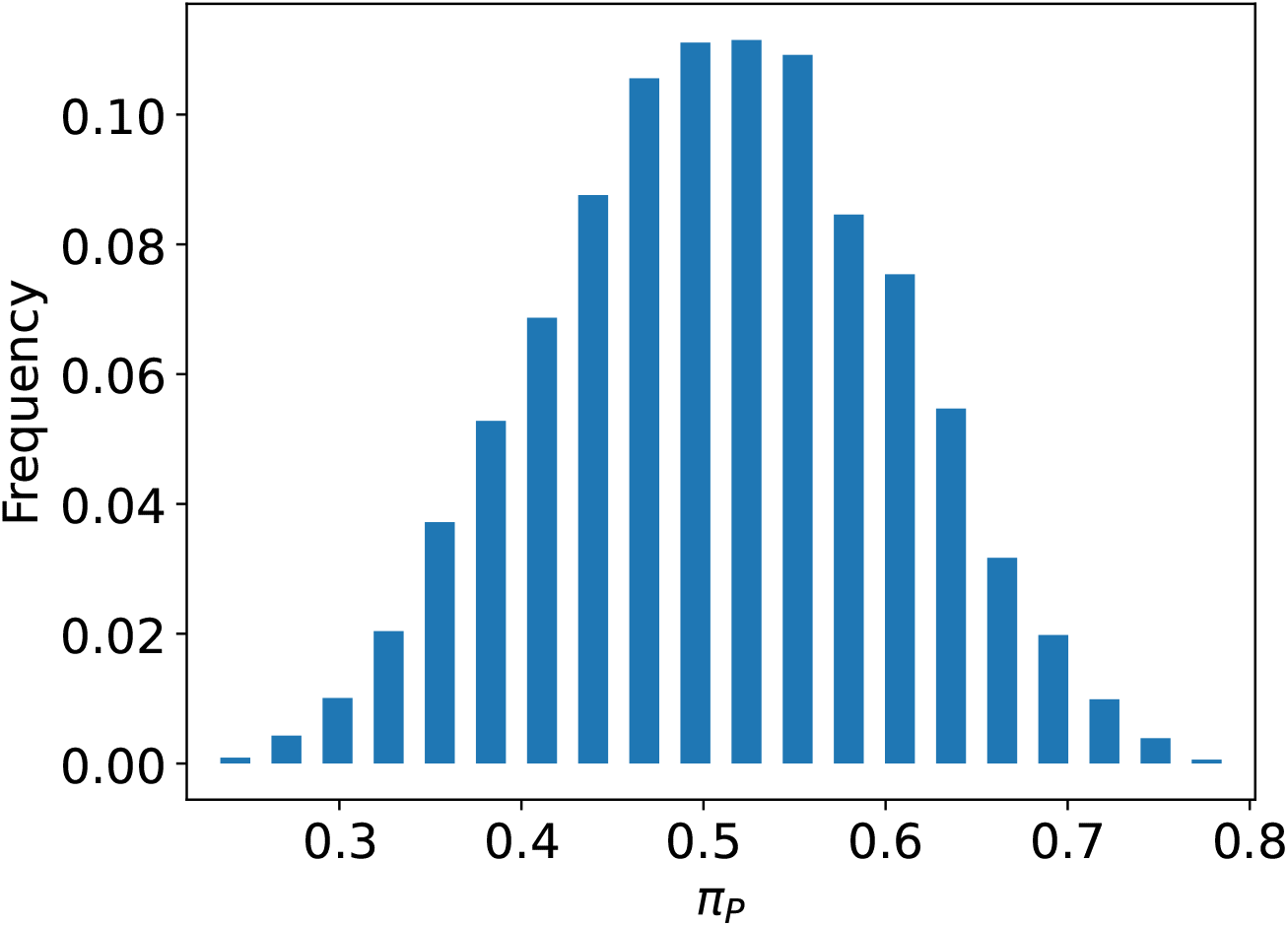
The distribution of the protein synthesis proteomic fraction *π*_*P*_ for random samplings the of λ_*i*_ parameters. Note the variability in *π*_*P*_ is commensurate with the input variability in the kinetics parameters, between 1*/*2 and 2 times the typical value.

### 4.3 Impact of molecular crowding

Besides calculating proteomic fractions for typical values of the enzyme parameters, we use our model to investigate how the proteomic fractions depend on key model parameters. The first question we address is if the macromolecular crowding in the cell restricts proteomic fraction in the cell. To do this we investigate the dependency of the model predictions on the macromolecular volume fraction *ϕ* (Fig. 2A-F, *ϕ* on the lower x-axis). First we confirmed that the parameter *κ* stays negative for all tested parameter values of *ϕ* (Fig. 2A). Second, we noted significant changes in the dilute limit *ϕ* → 0. When *ϕ* → 0 the ribosome and RNA polymerase proteomic fractions get close to zero (Fig. 2B, C). In contrast, the proteomic fractions associated with membrane synthesis and energy generation increase significantly when *ϕ* → 0. Therefore a cell with a low macromolecular fraction *ϕ* has a clear growth disadvantage. On the other hand, the protein and nucleotide synthesis proteome fractions saturate to their maximum predicted value when *ϕ* > 0.05 (Fig. 2B,C), indicating there is no further selection beyond beyond macromolecular volume fractions of about 0.05.

**Figure 2:**
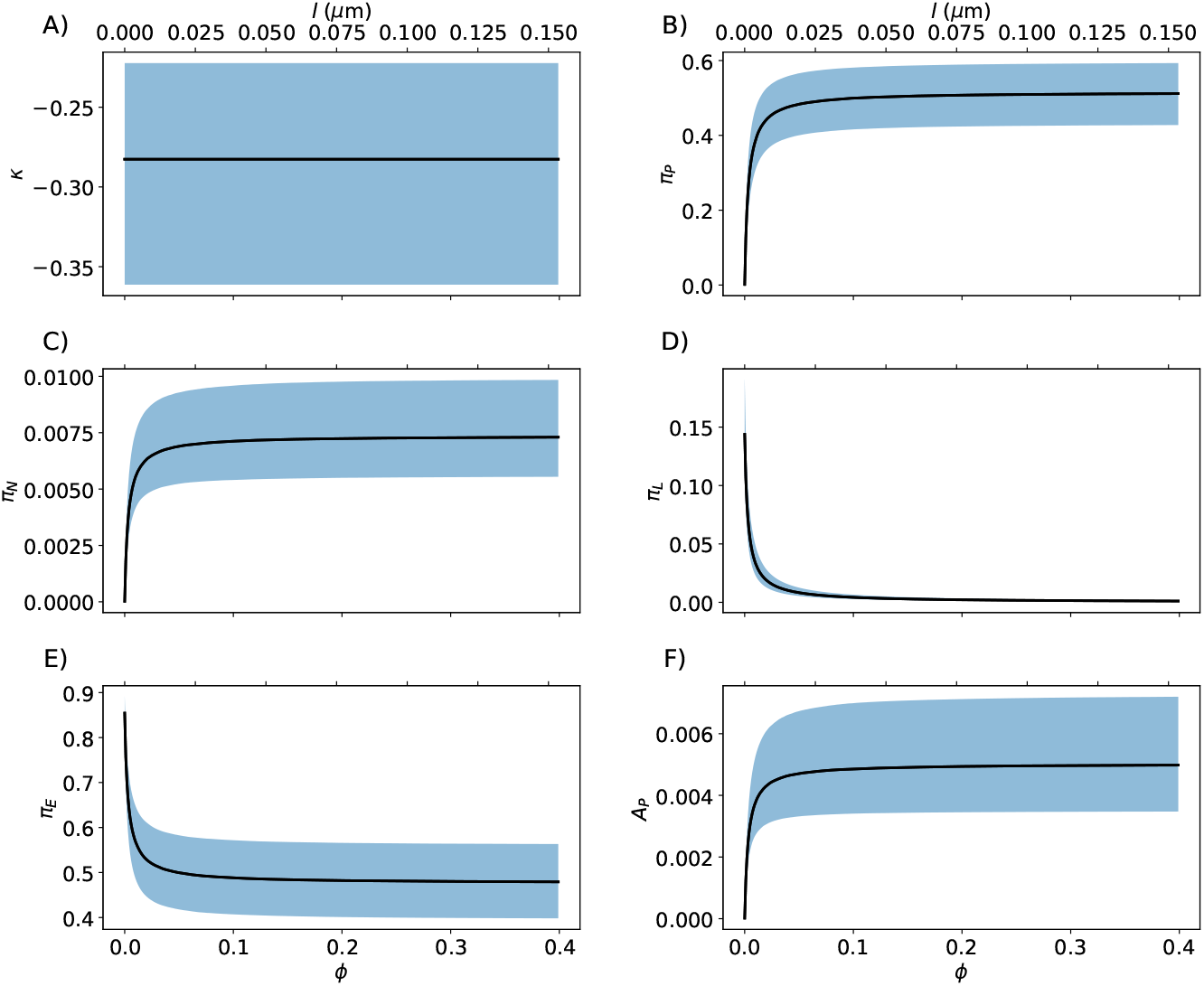
Functional dependency with the macromolecular volume fraction *ϕ* (lower X-axis) or volume to area ratio *l* (top X-axis) when assuming energy generation via glucose fermentation to acetate. The solid line is the median and the shaded are the confidence interval between the 0.2 and 0.8 quantiles. The dependent variables are previously defined protein fractions, *κ* and the fraction of the membrane area that is occupied by proteins *A*_*P*_, defined in (18).

Next we focus on the cell membrane area fraction *A*_*P*_ occupied by proteins, as defined in equation (18). The behavior of this quantity with the macromolecular fraction *ϕ* is shown in Fig. 2F. In this context *A*_*P*_ is small. The membrane synthesis pathway is the only component allocating a significant proteome fraction to the cell membrane and *π*_*L*_ is small for *ϕ* > 0.05 (Fig. 2D). However, as noted below this behavior changes when energy is generated by oxidative phosphorylation and the oxidative phosphorylation localizes to the cell membrane.

### 4.4 Impact of cell size

Since the volume-to-area ratio *l* is changing with the size of the cell and is different for cells of different shapes, we also investigate the dependency of the reduced model predictions with the volume to area ratio *l*. From equation (15) it follows that the model dependency on the parameter *l* is exactly the same as on the parameter *ϕ* (Fig. 2A-F, top -axis). Therefore *l* → 0 corresponds with the dilute limit described above. Indeed, as *l* → 0 the volume to area ratio is getting smaller and the cell is shifting most of its resources to membrane synthesis.

Our results suggest that there is a minimum volume to area size requirement to reach high *π*_*P*_ values, but above ∼ 0.01 *µ*m the maximal protein synthesis proteomic fraction is attained. That may sound counterintuitive but it is what may have been expected. If the volume to area is too small, then there will too few ribosomes to sustain the synthesis of enzymes in the membrane synthesis pathway. For the reference, the reported volume to area ratio for *E. coli* are in the range of 0.1 − 0.3*µ*m [33].

### 4.5 Energy generation via oxidative phosphorylation

We can repeat the simulations assuming instead that energy is generated via full oxidation of glucose, via glycolysis, the TCA cycle and oxidative phosphorylation. Unfortunately, we don’t have any estimate of the effective lambda for the complete oxidation of glucose. Here we will assume the same value of glucose fermentation to acetate (Table 1). However, we do know the ATP yield per molecule of glucose. We assume that most of the pathway localizes to the cell membrane, as it is the case in *E. coli*. This is clearly an approximation since during glycolysis some TCA steps take place in the cytoplasm. Here again *κ* < 0 (Fig. 3A) and the functional dependency of the proteomic fractions is similar to what observed with glucose fermentation as the energy source (Fig. 3B-E vs 2B-E). In contrast, when using oxidative phosphorylation there is a significant increase in the cell membrane area ratio occupied by proteins (Fig. 2F). This is the consequence of utilizing oxidative phosphorylation that localizes to the cell membrane. In this scenario volume and membrane crowding increase together.

**Figure 3:**
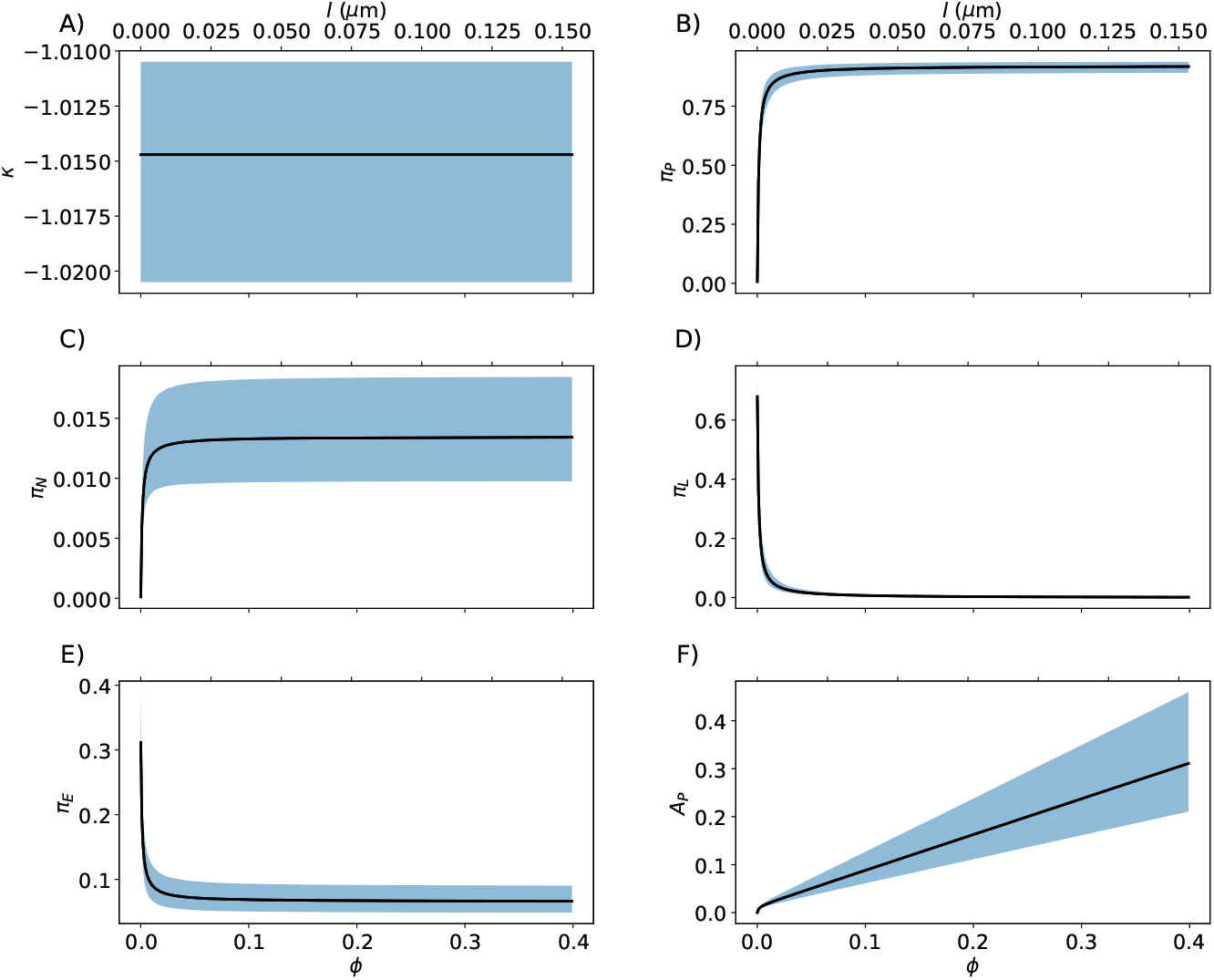
Functional dependency with the macromolecular volume fraction *ϕ* (lower X-axis) or volume to area ratio *l* (top X-axis), when assuming energy generation via complete oxidation of glucose (glycolysis + TCA cycle + oxidative phosphorylation). The solid line is the median and the shaded are the confidence interval between the 0.2 and 0.8 quantiles.

### 4.6 Impact of nutrient availability

So far we have assumed that the nutrients of the reduced model, glucose and amino acids, are in excess. Now we investigate the low glucose limit of *E. coli* cells growing on glucose as the only carbon source. To simulate glucose availability we set the λ of the glucose uptake step of glycolysis to λ_*G*_([Glucose]) = λ_*G*_[Glucose]*/*(*K* +[Glucose]), where [Glucose] denotes the glucose concentration and *K* the half-saturation constant for glucose uptake. We need to propagate this change to all pathways *i* = 𝒫, 𝒩, ℒ, ℰ. Since the λ of a pathway is the geometric mean of the λ of the individual pathway steps, the updated λ is given by

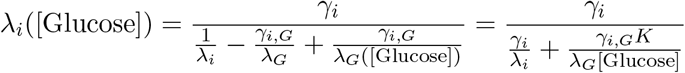

for *i* = 𝒫, 𝒩, ℒ, ℰ, where *γ*_*i,G*_ = 1*/*2 for *i* = 𝒫, 𝒩, ℒ and *γ*_*E,G*_ = 1.

We simulated the reduced model metabolism as a function of the glucose concentration, denoted by [Glucose], using as input the λ_*G*_ of the *E. coli* PTS system for glucose uptake and the extracellular glucose half-saturation constant *K*_*M*_ = 20*µM* [34]. We find that *κ* < 0 for all glucose concentrations tested (Fig. 4A), and therefore *π*_*P*_ = *π*_*P*,−_ is the only biologically relevant solution as reported above. With increasing [Glucose], the model predicts a small increase of the protein synthesis proteomic fraction *π*_*P*_, a decline of the proteomic fraction of the energy generating pathway (Fig. 4E) and an overall decrease of the proteome to lipid fraction making the cell membrane area (Fig. 4F). The decline of *π*_*E*_ with increasing [Glucose] can be explained by a reduction in the amount of glucose transporter required to sustain the glucose uptake with increasing the glucose concentration in the extracellular media. The increase in *π*_*L*_ with increasing [Glucose] is explained by the reduction in the glucose transporter proteomic fraction, which localizes to the cell membrane. The latter needs to be replaced by membrane lipids thus increasing the proteomic fraction of the membrane synthesis pathway.

**Figure 4:**
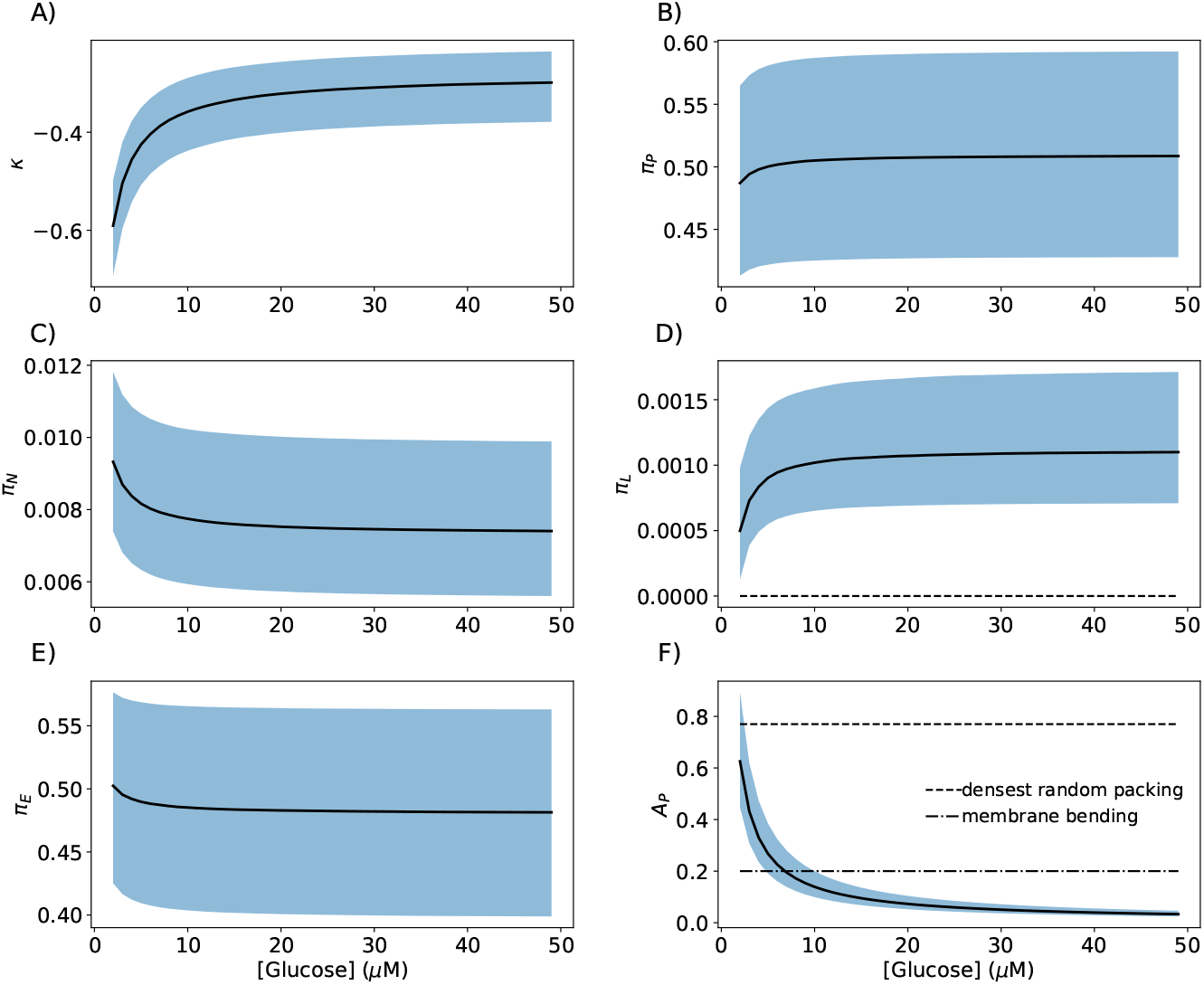
Functional dependency with the glucose concentration. The solid line is the median and the shaded are the confidence interval between the 0.2 and 0.8 quantiles.

For the smallest glucose concentrations the reduced model yields solutions with *π*_*L*_ ≈ 0 and *A*_*P*_ ≈ 1 (Fig. 4D). Based on packing constraints, *A*_*P*_ should not exceed the densest packing of disks (≈ 0.9 [35]) (Fig. 4F). In fact, we should not expect the perfect hexagonal configuration of the densest packing. At most random dense packings, which achieve a packing fraction of 0.77 [36]. Biophysical factors associated with membrane crowding are relevant as well. The cell membrane bends when the protein occupancy exceeds 20% [37], affecting cell viability in ways not accounted by our metabolic model.

Recall that in this simulation we have assumed a fixed cell geometry, *i*.*e*. fixed value of 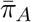. However, the cell could adjust its geometry or macromolecular volume fraction to increase 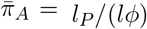, resulting in a feasible growth solution with *π*_*L*_ > 0 and *A*_*P*_ < *A*_*P*,max_, where *A*_*P*,max_ is some biologically relevant upper bound of the maximum membrane area fraction occupied by proteins. The cell can achieve that by reducing the volume to area ratio *l* and concomitantly decreasing the macromolecular content per cell to keep the macromolecular fraction *ϕ* constant. That observation is in agreement with empirical evidence. Bacteria and eukaryote cells have signaling pathways to reduce the cell size when nutrients are scarce. The proteogeometric constraint thus provides a hypothesis for the need of such adaptation.

## 5 Eukaryote cells

The extension of the proteogeometric constraint to eukaryote cells is not straightforward. Eukaryote cells contain organelles with their own membranes and so the cell contains membranes encapsulating each set of organelles and the cell membrane separating all the cell content from the extracellular media. In a first approximation, we can assume that the macromolecular density is homogeneous across compartments (cytosol, nucleus, mitochondria, etc). Under this approximation the volume balance in equation (4) can be used for eukaryote cells. The equation (5) for the cell membrane area balance remains valid as well, with the interpretation of *L* as the lipid content of the cell membrane and *E*_*L*_ as the amount of enzyme associated with synthesis of cell membrane lipids (not including the synthesis if intracellular membranes). In that context all equations (1)-(5) are valid and the proteogeometric constraint in equation (14) is derived.

Proteogeometric constraints are also valid for each organelle separately. The derivation for each organelle follows the same steps as we have done for the whole cell. To this end we denote by *π*_*o*_ and *π*_*Ao*_ the proteomic fractions localizing to the organelle *o* as a whole and to the organelle *o* membrane, respectively. We also denote by *π*_*Lo*_ the proteomic fraction associated with organelle membrane lipids synthesis that resides within the organelle. Finally, we replace *P* by *π*_*o*_*P* in the right hand side of the volume balance equation (4). With this definitions, and following the same steps as for the whole cell, we obtain the organelle specific proteogeometric constraints

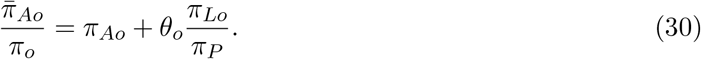

The investigation of these constraints in specific examples and, in particular, their coupling, will be the subject of future work.

## 6 Discussion

We have presented evidence demonstrating that the cell volume to area balance constrains cell metabolism beyond what is already accounted by proteomic balance constraints. Deviation of protein synthesis resources to the synthesis of cell membrane components implies a metabolic cost, which indirectly selects for compact cells with molecular crowding.

We may ask why flux balance models with a proteomic balance constraint alone, but no accounting of the volume to area constraint, have been successful in fitting experimental data. The answer is that usually such models use macromolecular composition of cells as an input and/or constraints. By doing so the models are restricting the analysis to the experimentally observed macromolecular proportions. In particular, the proportion between membrane lipids and enzyme proteins is usually assumed, rather than it emerges naturally from the model.

In a deeper sense, the challenge for systems biology is to make no assumptions about macro-molecular composition. With composition of available nutrients as the only input, the challenge is to predict what is the fastest growth in that specified environment, and what macromolecular composition supports such growth. We present evidence in this paper that such models need to account for cost of membrane biosynthesis, and for relationship between surface area and volume of the cell.

In conclusion, the proteogeometric constraint 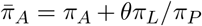 relates the cell geometry with its macromolecular composition.

## Supporting information

Supplementary Tables reporting parameters estimates.

## Supplementary Information

Tables with the reduced model parameter estimates.

## Data availability

All reported data is contained in the manuscript or the supplementary information. The reduced model and code to generate the figures are available at the github repository proteogeometric.

